# Temperature-dependant benefits and costs of cytoplasmic male sterility in snail *Physa acuta*

**DOI:** 10.1101/2024.07.23.604755

**Authors:** Sophie Bererd, Patrice David, Loïc Teulier, Sandrine Plénet, Emilien Luquet

## Abstract

Cytoplasmic male sterility (CMS) originates from a mito-nuclear conflict where mitochondrial genes induce male sterility and nuclear genes restore male fertility in hermaphrodites. The first observation of CMS in animals was reported recently in the freshwater snail *Physa acuta* where it is associated with two extremes divergent mitotypes D and K. The D individuals are male-sterile while male-fertility is restored by nuclear genes in K and are found mixed with the most common male-fertile N mitotype in natural populations (i.e., gynodioecy). We compared male and female fitness, growth rate, and metabolism between the three mitotypes at two temperatures as this factor influences CMS in gynodioecious plants via alteration of mitochondrial functioning. Temperature did not affect male-sterility which depended only on the mitotype and the presence of restorers. Our results provided evidence that CMS is beneficial to female fitness in the absence of restorers, while it is costly in their presence, fulfilling a key theoretical condition for the long-term maintenance of gynodioecy. Fitness benefits and costs are mediated by differences in body mass and enhanced at cold temperature suggesting that the system dynamics may vary according to thermal conditions in nature.

## INTRODUCTION

Opposed evolutionary interest between genomes can lead to conflicts between individuals (*e.g.* sexual conflict or parental-maternal, parents-offspring, social conflict) but also within individuals [1,2]. Intra-individual conflicts can occur between genomic compartments, such as nuclear and mitochondrial genes, which follow different strategies to maximize their own transmission [1]. Unlike nuclear genes transmitted by both parents, mitochondrial ones are inherited by only one parent, usually the female [3]. This disparity may lead to conflict over sexual reproduction in hermaphrodites [4,5] with the emergence of selfish mitochondrial lineages that sterilize the male reproductive function: the so-called cytoplasmic male sterility (CMS; [5,6]). The individuals carrying CMS are functionally female and are named male-steriles. In Angiosperms, where CMS is widespread, numerous examples highlight the association between divergent mitochondrial lineages and the failure to produce pollen [6–9]; such male-sterility would be caused by a dysfunction in the respiratory mitochondrial function [10]. As the sterilization of the male function is a loss of fitness for nuclear genes, the latter can engage in evolutionary arms race with mitochondrial genes [5,11,12]. Indeed, some nuclear genes can be selected to phenotypically suppress CMS [6,9,13,14]. These nuclear genes are commonly called restorers of fertility and are well described in many plant species [6,9,13,14]. Individuals carrying CMS and restoration genes are called restored hermaphrodites to distinguish them from normal hermaphrodites (*i.e.* non-CMS hermaphrodites). In natural populations, CMS results in the coexistence of hermaphrodites and male-sterile individuals, named gynodioecy [6,15]. The maintenance and dynamics of gynodioecy depends on how CMS and restorer genes can invade the populations. Firstly, theory predicts that for gynodioecy to be maintained CMS genes must benefit from a better transmission through a higher female fitness of male-sterile individuals called female advantage (FA; [6,16,17]). Meta-analyses show empiric evidence for a FA in a majority of investigated gynodioecious plant species although its magnitude, and the life stage at which it is expressed (*e.g.,* number of seeds or germination rate), vary greatly among species [16,18]. For example, FA consisted in an overproduction of seeds in male-sterile individuals of *Silene acaulis* and *Echium vulgare* [18]. However, fertility-based female advantage would be in balance with the cost of carrying/expressing CMS genes [19–21]. For example, in *Plantago maritima* [22], females (male-steriles) produce more seeds (FA) but they have a lower germination rate than those of hermaphrodites, which can be interpreted as CMS cost as suggested by Dufaÿ et al [19]. Although such a CMS cost has been poorly investigated empirically, its expression is probably crucial in the presence of restorer genes; when restorers are abundant, CMS is carried mostly by restored hermaphrodites, and the latter may go to fixation if their female fitness is not inferior to that of normal hermaphrodites (not carrying CMS). The system would then revert back to a 100% hermaphroditic population. However few studies compare female fitness of restored versus normal hermaphrodites and interpret the difference as a cost of CMS [21]. Another condition for the maintenance of gynodioecy is that restorer alleles do not go to fixation, implying that they also incur a fitness cost (expressed on male and/or female fitness; [6,8,13,19]). The existence of such a cost has been reported in restored hermaphrodites of *Lobelia siphilitica* (lower pollen viability; [23]) or *Brassica napus* (reduced flower size, stamen length and pollen counts depending on nuclear-cytoplasmic assembly; [24]). Consequently, the changes in frequency of CMS and restorer genes driving the dynamics of gynodioecious populations (loss/fixation of CMS and restorers genes, frequency-dependent oscillations) depend on the magnitude of both the FA and costs (CMS and restoration; [19]). However, in practice, the cost of CMS and the cost of restoration are difficult to disentangle/distinguish because these can act on the same fitness traits and it also requires comparing the fitness of individuals carrying or not CMS and restorer genes, which is often impossible.

Environmental conditions may also influence the dynamic of gynodioecy in natural populations. Since CMS is associated with mitochondria [10], which plays a vital role in metabolism, it may be sensitive to factors affecting mitochondrial functioning such as stress or temperature. Firstly, environment may influence the effects of CMS genes on male sterility [25,26]. In some plants, male fertility of individuals with CMS genes can be totally or partially restored by environmental factors such as humidity [27] or temperature [28,29] without the intervention of restorer genes. For example, some authors reported a spontaneous restoration of male fertility in rape [30] and in barley [31,32] by a reactivation of pollen growth in male-sterile individuals (*i.e.* bearer of CMS genes) at high temperature, while others observed an increase of pollen production at low temperature in male-steriles in maize [28,29]. Secondly, environmental conditions may also influence the effects of restorer genes on male fertility [33]. For example, the restoration of fertility in chieves is triggered only at high temperature because some specific nuclear restorer genes are temperature-sensitive [33,34]. As a consequence, CMS and restorer genes may become neutral in certain environmental conditions and thus may be cryptically maintained in some populations. Thirdly, environmental conditions may also influence FA and costs of CMS and restoration [16]. However, direct tests of environmental effects on FA and costs are scarce and show mixed results: female fitness was affected by soil moisture is *Nemophila menziesii* [35], but no effect on the magnitude of FA and cost was detected for temperature in *L. siphilitica* [36] and for light availability in *Geranium sylvaticum* [37]. In addition, the observations of variation in FA and costs among populations could result from such effects driven by environmental differences and explain different dynamics of gynodioecy [38,39]. Higher female frequency is found in stressful environment / poor quality habitats [40], which suggest that these environment could increase the magnitude of both FA for male-steriles and costs for hermaphrodites by affecting their fitness [36].

Consequently, understanding the evolutionary dynamics of gynodioecious systems is complex as it is challenging to evaluate the FA and costs (CMS and restoration) and their potential environmental dependency. A relevant biological model to take on this challenge has been recently discovered by David et al [41]. They reported the first observation of animal CMS and gynodioecy in *Physa acuta* (Physidae, Hygrophila, Gastropoda), a freshwater snail with a worldwide geographical distribution [42,43]. It is a simultaneous hermaphrodite with preferential outcrossing, *i.e.* eggs are self-fertilized only when no mate is available to provide allosperm [44–46]. However, some individuals were found to be unable to sire offspring. This male sterility is associated with a mitochondrial genome – called mitotype D – characterized by an extreme divergence compared to the classically described mitochondrial genome – called mitotype N (median divergence of 44.4% for nucleotides; [41]). Interestingly, a third mitochondrial lineage – called mitotype K – reported lately is also extremely divergent from both D and N lineages (DNA divergence between K and N ranged from 35% to 57%, and between K and D ranged from 41% to 61%; [47]). While they harbour a hermaphrodite phenotype in their natural population, K individuals become male-sterile after the introgression of K mitogenome into a naïve nuclear genomic background, revealing for the first time in animals the existence of nuclear restorer genes and a cryptic CMS system [47], *i.e.* the (hermaphroditic) K individuals in their native population are in fact restored hermaphrodites.

The aim of the present study is to investigate how the environment influences the key parameters of the dynamics of gynodioecy (*i.e.,* male fitness of CMS gene bearer with or without restorer genes, FA and CMS and/or restoration costs). We focused on temperature as it may affect overall life history traits [48,49], through a strong influence on mitochondrial functioning; and because it is known to restore male fertility in CMS plants [28,30–32] and reverse male fertility (neutralization of restorer gene effects) in restored hermaphrodites [33,34] in plants. We raised the three *P. acuta* mitotypes (N, D and K) in two thermal conditions, low (18°C) and high (25°C) temperatures to compare male fitness (male sterility, seminal vesicle area and number of spermatozoids) and female fitness (eggs production), growth rate and *in vivo* metabolic rate. These temperatures are in the wide range occupied by *P. acuta* (the highest reproduction rate is between 15°C and 25°C; [50]). At the time when the experiment was set up, only individuals with the genetic background of their native population were available, so that (i) D individuals can be expected to be non-restored male-sterile individuals (restoration has never been observed for the D CMS mitotype); (ii) K individuals are expected to be restored hermaphrodites; (iii) N individuals, from the same origin, are non-CMS individuals but expected to mostly carry K-specific restorer alleles in their nuclear genome, as they share the same background as their K congeners from the same population. Firstly, we tested if temperature affects the effect of CMS and restorer genes on male fitness by comparing individuals carrying CMS (D) to their non-CMS counterparts (N) in the absence of D-specific restorers; and by doing a similar comparison between K and N, this time in the presence of K-specific restorer genes, respectively. Secondly, we measured benefits and costs of CMS on the female function, both in non-restored and restored contexts, and how they are affected by temperature.

## MATERIAL AND METHODS

### Experimental design

We used three laboratory *P. acuta* lineages: mitotype N hermaphrodites, mitotype D male-steriles and mitotype K restored hermaphrodites. They have been obtained from wild populations near Lyon (southeastern France; Table S1; [41,47]) and raised in laboratory at 25°C for 13 generations for N and K and 26 generations for D. Originally, N and K snails come from the same two populations (Table S1); they have been separated at the first generation and have since been reproduced separately. The D snails (male-steriles) are fertilized by N snails through pair-crosses at each generation to obtain D offspring. The three mitotypes therefore share the same nuclear background. During the experiment, all the snails were reared individually in small boxes (90 ml) with dechlorinated water changed once a week and were fed with *ad libitum* boiled and mixed lettuce twice a week. The experiment took place in two temperature-controlled rooms, with two defined constant temperatures, a low and high temperatures (respectively, 18.29 ± 0.15°C and 25.7 ± 0.30°C), with a 12h light-dark photoperiod. At 11 days post-hatching, 100 snails per mitotype were randomly assigned to one of the two thermal treatments (50 individuals / mitotype / temperature, total n = 300 individuals).

### Life-history traits measurements

#### Male and female fitness

##### Male sterility

To test male sterility of the three mitotypes, we assumed that each focal individual was mature with a minimum weight of 0.04g (45 days after the start of the experiment at 18°C and 22 at 25°C; respectively, 56 and 33 days post-hatching). We paired each focal individual with a virgin albino for copulation for 24 hours. Each partner (focal and albino) was thereafter individualized for 3 days to lay eggs in small rearing boxes. The albino individuals are N mitotype, hermaphroditic snails (*i.e.* male-fertile and non-CMS), originating from the Montpellier Area (∼300km south of Lyon) and have been maintained as a large laboratory population for over 80 generations, and were used for male-fertility assessment in [41,47]. 5 days after laying at 25°C and 7 days after laying at 18°C, the pigmentation of the offspring produced by the albino partner was determined through the eye color of (still unhatched) embryos under a binocular microscope. If offspring from albino partner were pigmented, the focal individual was considered as male-fertile. On the contrary, snails were considered sterile when the albino partner laid no eggs or produced only albino offspring (*i.e.* results from a self-fertilization; see [41] for more details).

##### Male reproductive traits: seminal vesicle area and number of spermatozoids

After copulation and egg-laying, all focal individuals of the three mitotypes were dissected to extract the seminal vesicle, organ of sperm maturation and storage. Seminal vesicle area (mm²) was estimated by photographic analysis using the ImageJ software (Olympus SZX9, Olympus SC50 camera; [51]) to the nearest 0.001mm. Sperm counts were made on two independent 0.1 µl aliquots of each extract at 400X magnification (Keyence VHX-7000), using a hemocytometer after crushing the vesicle (see detailed protocol in [41]).

##### Female fitness: number of eggs

Female fitness of focal snails from the three mitotypes was estimated as the number of eggs laid after mating with albino individuals. Eggs were counted using a binocular microscope, 5 and 7 days after laying respectively for 25°C and 18°C.

#### Growth rate: whole body mass and maximal shell length

Snail mass and size were measured at sexual maturity, *i.e.* 22 days after the start of the treatment for individuals at 25°C (33 days old snails since hatching) and 45 days for those at 18°C (56 days old snails). Whole body mass (shell + body) was measured with a precision scale to the nearest 0.001 mg. Maximal shell length was assessed using photography analyzed with the software ImageJ [51] at the nearest 0.001mm. Growth rate was calculated by a ratio between whole body mass or maximal shell length and number of days since the start of the experiment.

#### *In vivo* metabolic rate

*In vivo* metabolic rate of all focal snails was estimated at both temperatures, 25°C and 18°C at respectively 28 and 53 days after the start of the experiment, through whole-organism oxygen consumption (MO2, in mgO2/h) measured by a stop-flow respirometry protocol during 90 to 120 min. Briefly, each individual was placed in a respirometric chamber filled with fully aerated water (V = 1mL) built with a 5mL seringe, connected to a peristaltic pump (Watson Marlow, model 205S), allowing continuous water circulation (flow rate: 7 ml/min) to homogenate oxygen concentration in the chamber. Oxygen concentration was recorded every minute by an optode (robust probe, Pyroscience, Germany) connected to a computer through a specific interface (Firesting Pyroscience, Germany). Metabolic chambers were immersed into a tank of water at 25°C or 18°C, to keep water temperature constant during all the experiment. MO2 of each snail was estimated following the equation: MO2 = (Δ[O_2_] x (Vch-Vsnail))/Δt), with [O2] the concentration of oxygen in the chamber (mgO_2_.mL^−1^), Vch the volume of the chamber (mL), Vsnail the volume of the snail (mL) and t the time (h).

### Statistical analysis

A generalized linear model (GLM) following a binomial distribution was used to test the influence of temperature and mitotype on male sterility by an analysis of variance (ANOVA). The influence of whole-body mass, temperature, mitotype and the interactions of these factors on the surface of the seminal vesicle was tested by an ANCOVA on the linear model. A GLMM following a quasi-Poisson distribution to consider overdispersion, was carried out to test by an ANOVA the influence of the vesicle surface, temperature, mitotype as well as their interactions on the number of spermatozoa. An individual random effect was used to consider 2 counts by individuals. A GLM following a quasi-Poisson distribution to consider overdispersion was used to test the influence of whole-body mass in addition to temperature, mitotype and their interactions on the number of eggs. A linear model was built to test the influence of temperature, mitotype and their interactions on growth rate using an ANOVA. *In vivo* metabolic rate was analysed using a linear model by an ANOVA to test the effect of mitotype, temperature and their interaction. All statistical analyses were conducted in RStudio version 4.3.2 [52]. Functions from packages stats, lme4 and emmeans were used. Figures were produced using ggplot2 and ggpbur packages. Post-hoc pairwise comparisons (using a Tukey test) were performed on simplified models (*e.g.* interactions removed when non-significant) to identify differences between mitotypes (according to temperature when the interaction was significant). Estimates (± SE) of the main factor temperature were indicated in the results considering low temperature (18°C) as the intercept.

## RESULTS

### Male sterility

Male sterility depended on mitotype and on temperature (interaction: χ²_2, 239_ = 1.97, p = 0.373; mitotype effect: χ²_2, 239_ = 121.64, p < 0.0001; temperature effect: χ²_2, 239_ = 10.45, p = 0.001). D individuals were almost all sterile whatever the temperature (Fig. 1). The proportion of male-fertility was much lower in D than in N and K (in logit scale, D - N = −4.71 ± 0.71, z = −6.68, p < 0.001; D - K = −5.46 ± 0.86, z = −6.31, p < 0.001), while N and K were not significantly different (in logit scale, N - K = 0.75 ± 0.88, z = 0.84, p = 0.67), and mostly male-fertile whatever the temperature (Fig. 1).

**Figure 1.**
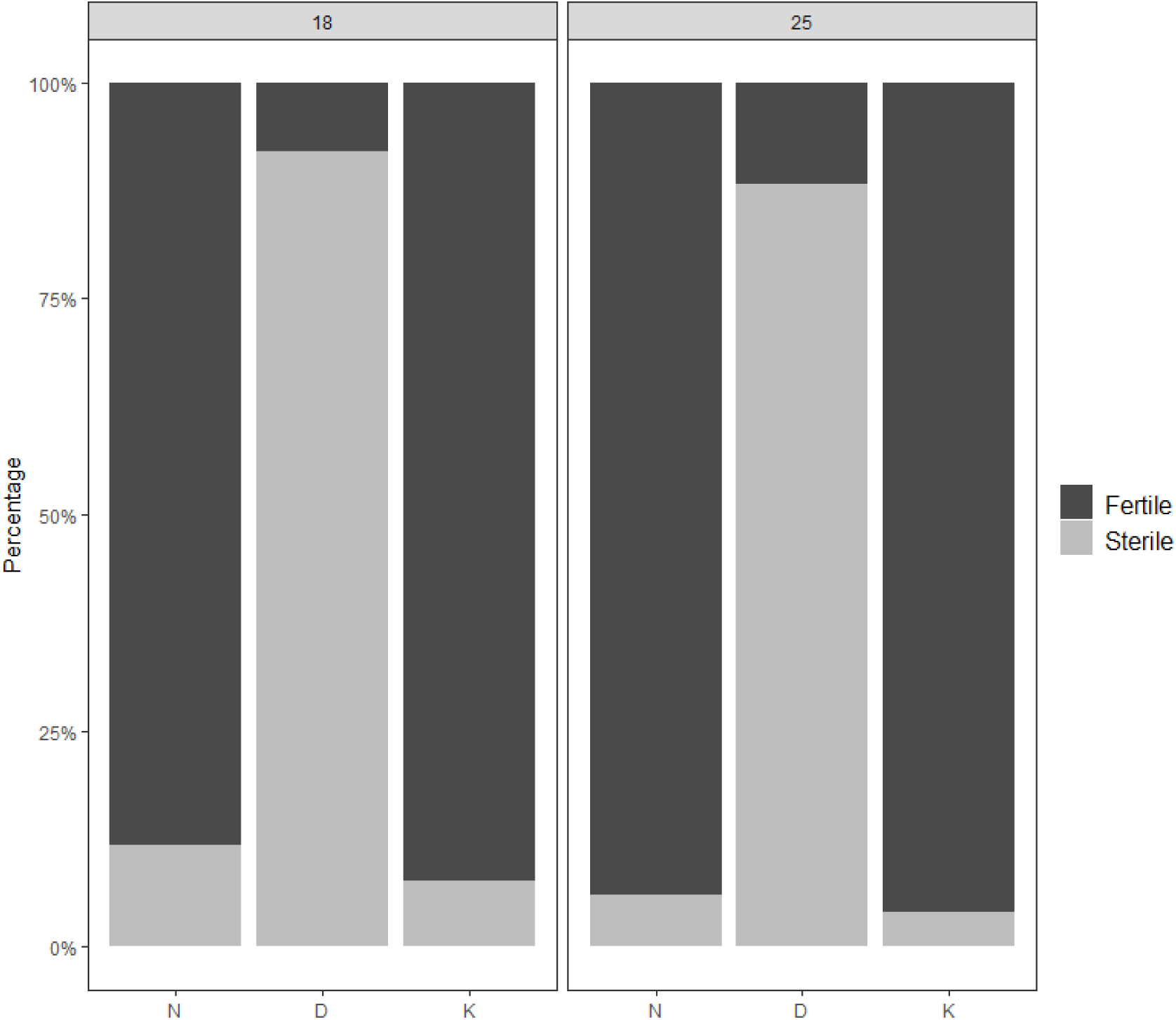
Percentage of male-fertile and male-sterile individuals in both N, D and K *P acuta* mitotypes at 18 and 25 °C. Total sample sizes were 51, 50 and 52 at 18°C and 50, 51 and 50 at 25°C for respectively N, D, and K.

As the number of N and K snails assigned male-sterile and D assigned male-fertile by the male sterility test is limited (at 18°C: 6 N and 4 K male-sterile, 4 D male-fertile; at 25°C: 3 N and 2 K male-sterile, 6 D male-fertile), no statistical analysis has been carried out and the following statistical analyses were conducted only on N and K male-fertile individuals (*i.e.* normal hermaphrodites and restored hermaphrodites respectively) and D male-sterile individuals. In the supplementary section, we added figures allowing to understand whether these individuals are false-positive or false-negative (see supplementary Fig. S6 and S7) and to discuss about incomplete CMS penetrance.

### *In vivo* metabolic rate (MO2)

MO2 was not influenced by mitotype (F_2, 250_ = 0.0456, p = 0.9554) or temperature (F_1, 250_ = 1.4569, p = 0.2286) and interaction was not significant (F_2, 250_ = 0.5429, p = 0.5817; Fig. 2).

**Figure 2.**
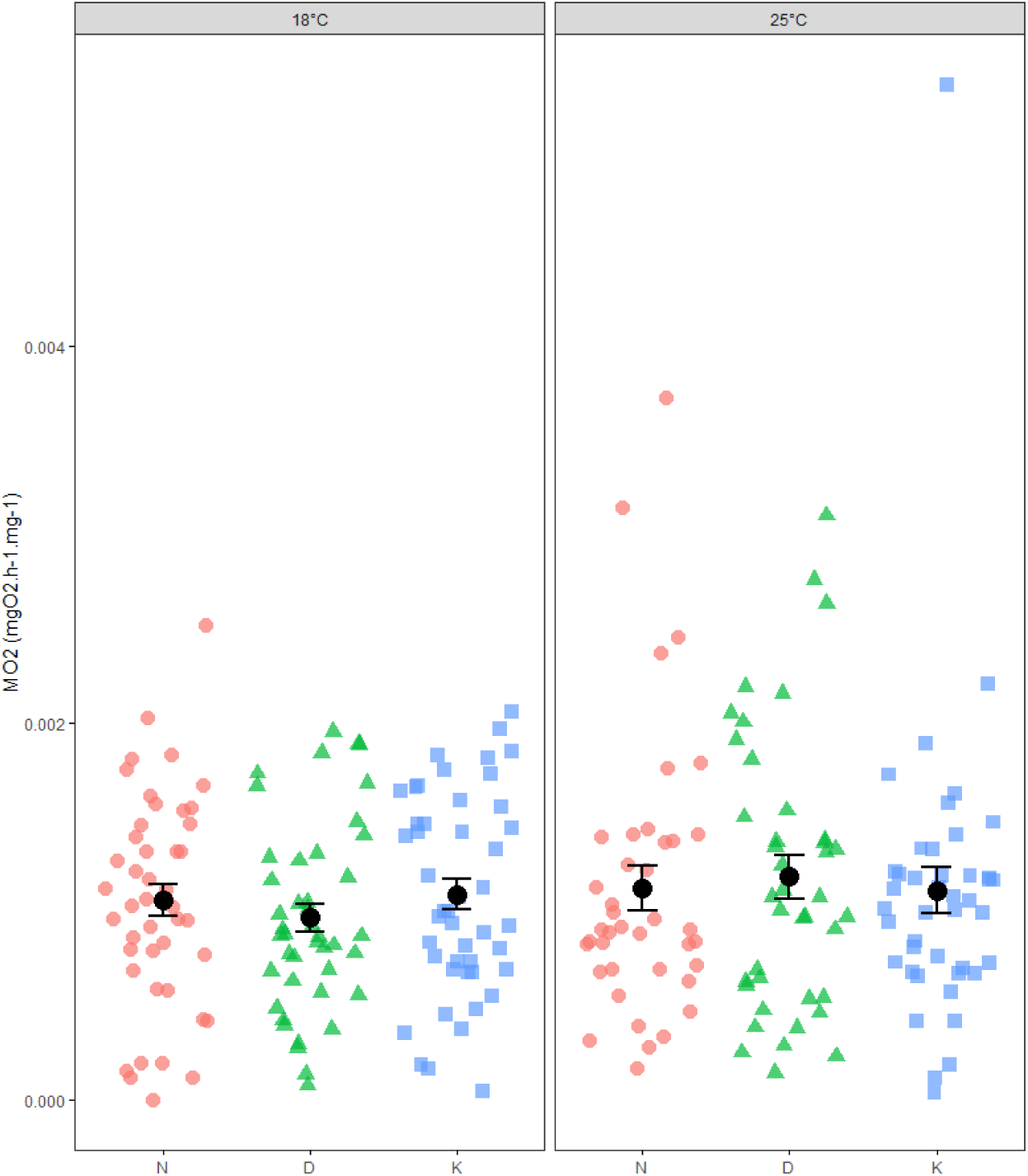
*In vivo* metabolic rate (MO2, in mgO_2_.h^−1^.mg snail^−1^) for normal *P. acuta* hermaphrodites N (red circle), male-sterile D (green triangle) and restored hermaphrodites K (blue square) at 18 and 25°C. Dots correspond to observed values.

### Growth rate

#### Body mass

Growth rate from the body mass (BM growth) depended on the interaction between mitotype and temperature (interaction: F_2,272_ = 4.011, p = 0.019; mitotype effect: F_2, 272_ = 21.67, p < 0.001; temperature effect: 1.229e-03 ± 1.153e-04, F_1, 272_ = 373.33, p < 0.001; Fig. 3A, see Fig. S2 for raw BM).

**Figure 3.**
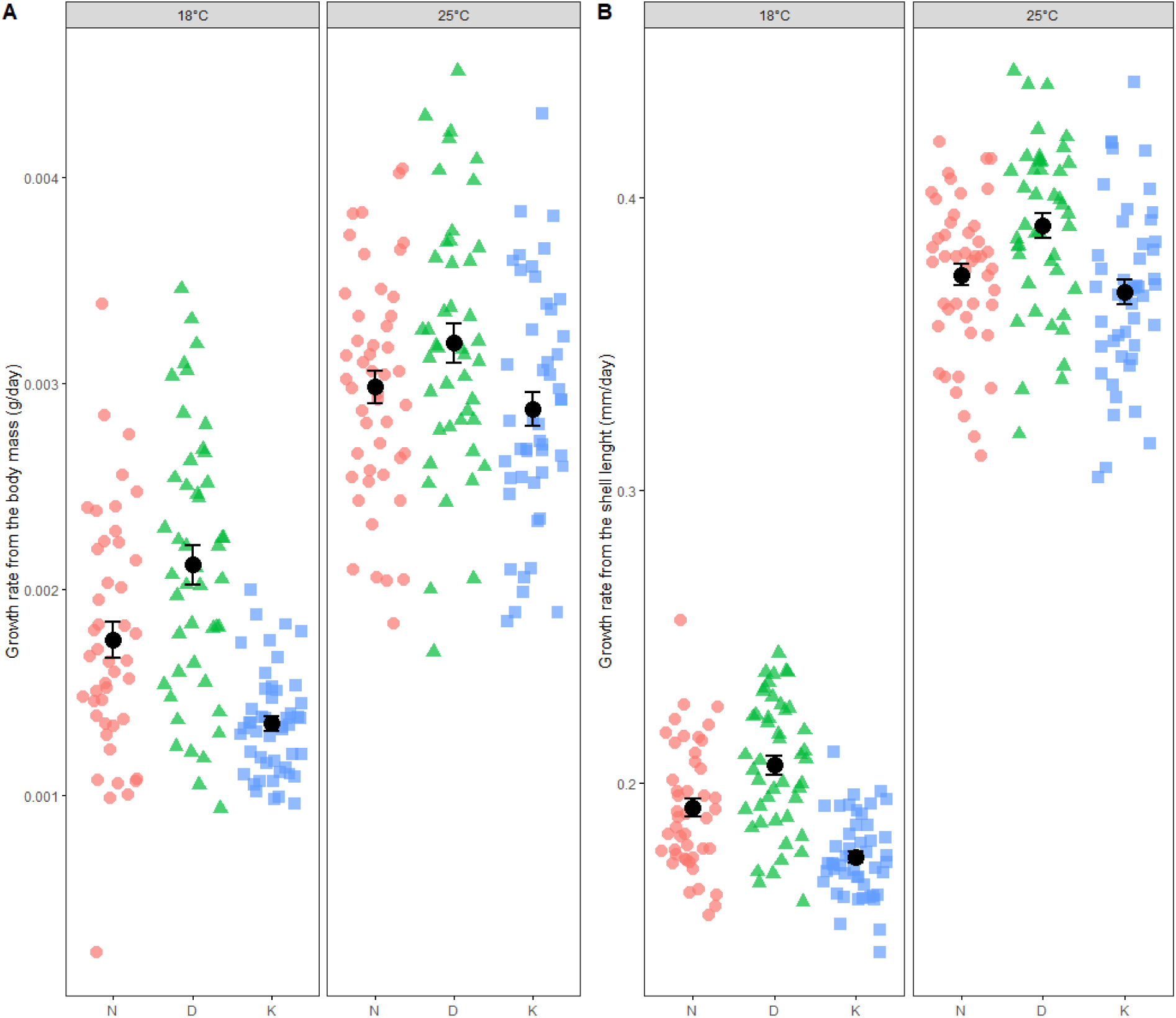
**A -** Growth rate from the body mass (BM) for normal *P. acuta* hermaphrodites N, male-sterile D and restored-hermaphrodites K at 18 and 25°C. **B -** Growth rate from the shell length for N, D and K mitotypes at 18 and 25°C.

At 18°C, the BM growth of the three mitotypes differed. The BM growth rate of D snails was 37% higher than those of K and 17% of N (D - K: Est.±SE = 0.0007 ± 0.0001, t_272_ = 6.76, p < 0.001; N - D: −0.0003 ± 0.0001, t_272_ = −3.16, p = 0.005; Fig. 3A). K snails had a significantly 23% lower growth than N ones (N - K = 0.0004 ± 0.0001, t_272_ = 3.53, p = 0.001; Fig. 3A).

At 25°C, the ranks of the three mitotypes with respect to mean body mass growth rate were K<N<D as they were at 18°C; however the differences were quantitatively smaller and significant only between D and K where K snails were 11% smaller than D (D - K: Est.±SE = 0.0003 ± 0.0001, t_272_ = 2.78, p = 0.016; N - K: 0.0001 ± 0.0001, t_272_ = 0.96, p = 0.604; N - D: 0.0002 ± 0.0001, t_272_ = −1.83, p = 0.163; Fig. 3A).

#### Maximal shell length

The results on shell length growth rate were similar to those obtained with body mass, with the ranks K<N<D at both 18°C and 25°C. However, the interaction between temperature and mitotype was not significant (F_2, 272_ = 1.37, p = 0.26), while mitotype (F_2, 272_ = 24.73, p < 0.005) and temperature effects were (faster growth at 25°C than 18°C: 0.18 ± 0.005, F_1, 272_ = 4110.65, p < 0.0001). All pairwise differences between the three mitotypes were highly significant (N – D: Est.±SE = −0.016 ± 0.004, t_272_ = −4.35, p = 0.0001; D - K: 0.027 ± 0.003, t_272_ = 7.59, p < 0.0001; N - K: 0.011 ± 0.003, t_272_ = 3.21, p = 0.004; Fig. 3B, see Fig. S3 for raw shell length). D snails were 4% longer than N and 8% than K and N were 5% longer than K.

Colored-shape dots correspond to observed values (red circles for N normal hermaphrodites, green triangle for D male-steriles and blue square for K restored hermaphrodites) and black dots are mean ± standard error. Recall that BM growth rate and length growth rate have been obtained by dividing mass or length by age allowing to compare between temperatures (measured was taken at 45 and 22 days for 18 and 25°C treatments, respectively). Graphs with raw body mass and raw length are given in supplementary figure S2 and S3, respectively.

### Male Fitness

#### Seminal vesicle area

The seminal vesicle area was influenced by the interaction between body mass and mitotype, the main mitotype effect and marginally by temperature (Table 1; Fig. 4A). The seminal vesicle area of D snails was much smaller, in the whole range of body masses, than those of N and K whatever the temperature (D - N: 1.20 ± 0.064, t_264_ = 19.07, p < 0.001; D - K: −1.146 ± 0.073, t_264_ = −16, p < 0.001; Fig. 4A). The vesicle area did not differ between N and K (0.054 ± 0.071, t_264_ = 0.77, p = 0.721; see Fig. 4A).

**Table 1.** Results of linear model ANCOVA on the effects of body mass, mitotype and temperature on seminal vesicle area.

| Effects | NumDF, DenDF | F | P |
| --- | --- | --- | --- |
| Body mass (BM) | 1, 259 | 27.162 | **<0.001** |
| Mitotype (M) | 2, 259 | 233.166 | **<0.001** |
| Temperature (T) | 1, 259 | 3.491 | 0.063 |
| BM x M | 2, 259 | 4.608 | **0.011** |
| BM x T | 1, 259 | 0.006 | 0.940 |
| M x T | 2, 259 | 0.715 | 0.490 |
| BM x M x T | 2, 259 | 1.170 | 0.312 |

**Figure 4.**
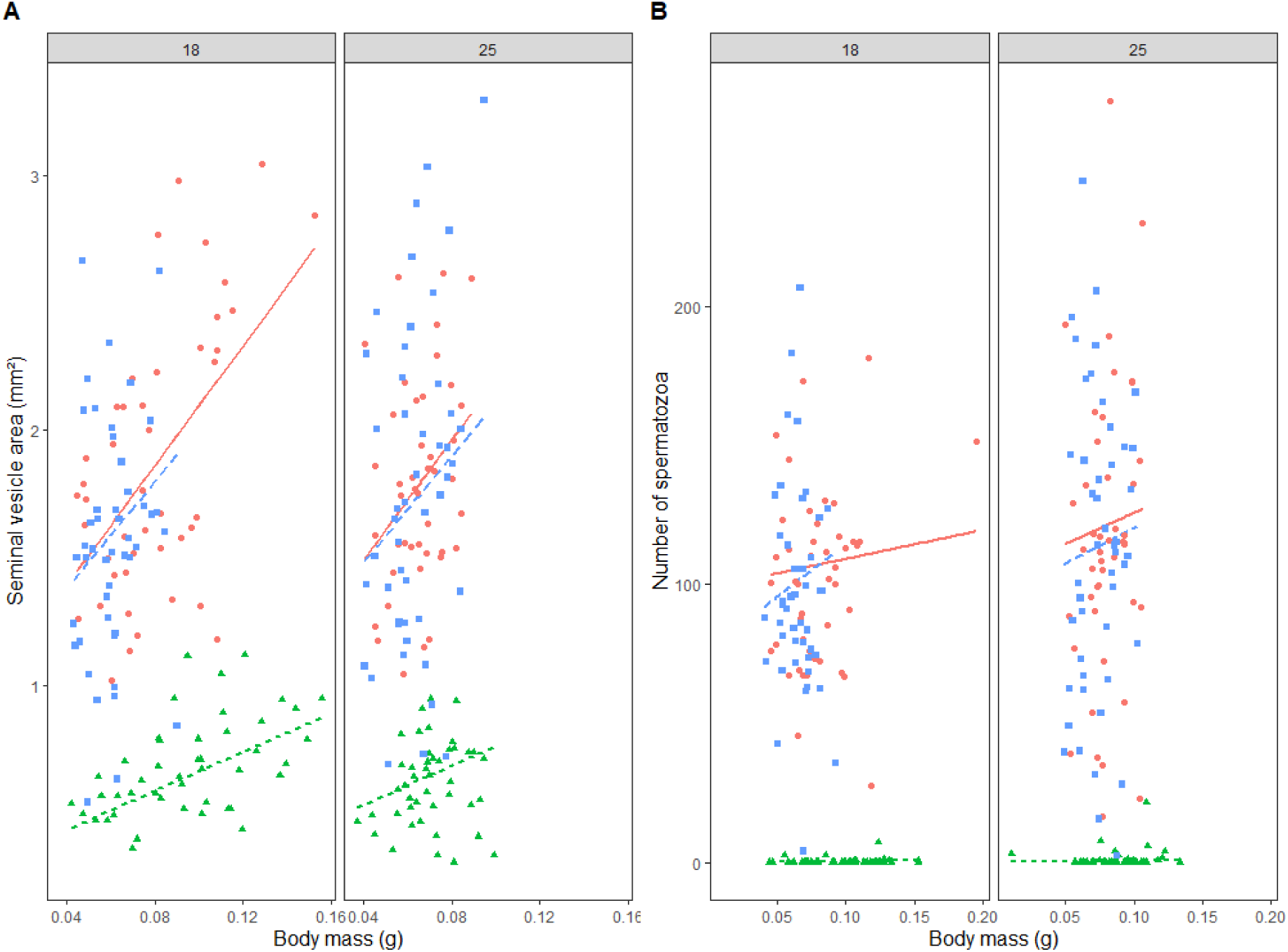
**A -** Seminal vesicle area according to body mass for normal *P. acuta* hermaphrodites N, male-sterile D and restored hermaphrodites K at 18 and 25°C. **B -** Number of spermatozoa (means between two individual counts for a sake of clarity) according to body mass for normal N, D and K mitotypes at 18 and 25°C. Raw distributions of sperm counts are shown in figure S4. Lines represent the model prediction and dots correspond to observed values (red circles and plain line for N normal hermaphrodites, green triangle and small dashed line for D male-steriles and blue square and large dashed line for K restored-hermaphrodites).

#### Number of spermatozoids

The number of spermatozoids was influenced by the mitotype and the temperature (more spermatozoids at 25°C than 18°C: 4.31 ± 0.19) and all interactions were not significant (Table 2; Fig. 4B). D snails produced practically no spermatozoa whatever the temperature (D - N: 5.42 ± 0.56, t_167_ = 9.60, p < 0.0001; D - K: −5.41. ± 0.57, t_167_ = −9.57, p < 0.0001; Fig. 4B). The number of spermatozoids did not differ between N and K (0.01 ± 0.06, t_167_ = 0.18, p = 0.982; Fig. 4B; Fig. S4).

**Table 2.** Results of linear model ANCOVA on the effects of body mass, mitotype and temperature on number of spermatozoa.

| Effects | NumDF, DenDF | Chisq | P |
| --- | --- | --- | --- |
| Body mass (BM) | 1,160 | 2.285 | 0.131 |
| Mitotype (M) | 2,160 | 100.701 | **<0.001** |
| Temperature (T) | 1,160 | 6.107 | **0.013** |
| BM x M | 2,160 | 5.249 | 0.072 |
| BM x T | 1,160 | 2.368 | 0.124 |
| M x T | 2,160 | 0.285 | 0.867 |
| BM x M x T | 2,160 | 2.067 | 0.356 |

### Female Fitness

#### Number of eggs

Egg production depended on mitotype (F_2,253_ = 8.98, p = 0.0001) and temperature (egg production higher at 18°C than 25°C: −0.22 ± 0.10, F_1, 253_ = 13.99, p < 0.001; interaction: F_2, 253_ = 1.53, p = 0.219). D snails produced more eggs than N and K snails whatever the temperature (N - D: Est.±SE = −0.19 ± 0.067 (log scale), z = −2.86, p = 0.012; D - K: −0.29 ± 0.071, z = 4.07, p = 0.0001; see Fig. 5A). Egg production did not differ significantly between N and K snails (N – K = 0.095 ± 0.073, z = 1.31, p = 0.391; Fig. 5A). Relative to N, female fertility of D is estimated to be 20% higher. while the (non-significant) decrease in fertility between K and N is −9%. The fertility rankings (D>N>=K) mirrored results obtained with body mass; when body mass was accounted for, mitotype and temperature no longer had significant effects on egg production, suggesting that differences depended on body mass (Table 3; Fig 5B; Fig. S5).

**Figure 5.**
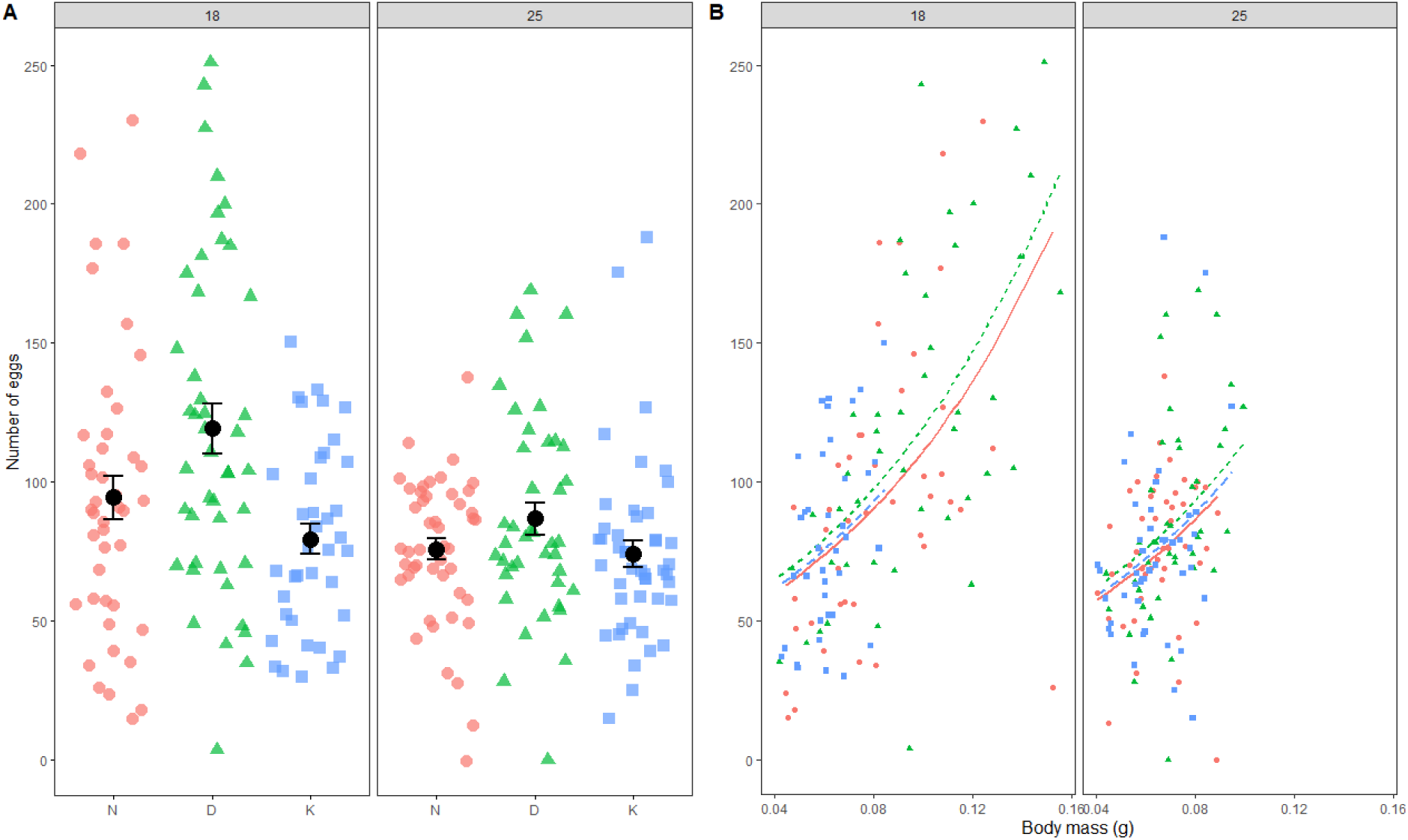
**A -** Number of eggs for *P. acuta* normal hermaphrodites N, male-steriles D and restored hermaphrodites K at 18 and 25°C. **B -** Number of eggs according to body mass for N, D and K mitotypes at 18 and 25°C. Lines represent the model prediction, colored shape dots correspond to observed values and black dots are mean ± standard error (N = red circle, plain line; D = green triangle, small dashed line; K = blue square, large dashed line). Raw distributions of egg number produced by focal individuals are shown in figure S5.

**Table 3.** Results of linear model ANCOVA on the effects of body mass, mitotype and temperature on number of eggs.

| Effects | NumDF, DenDF | F | P |
| --- | --- | --- | --- |
| Body mass (BM) | 1, 247 | 70.297 | **<0.001** |
| Mitotype (M) | 2, 247 | 0.767 | 0.465 |
| Temperature (T) | 1, 247 | 0.656 | 0.419 |
| BM x M | 2, 247 | 0.540 | 0.583 |
| BM x T | 1, 247 | 0.044 | 0.835 |
| M x T | 2, 247 | 0.295 | 0.745 |
| BM x M x T | 2, 247 | 1.065 | 0.346 |

## DISCUSSION

The aim of the present study was to investigate how temperature influenced the male fitness of CMS genes bearer without or with restorer genes (D and K mitotypes, respectively), the female advantage and costs (CMS and/or restoration), key parameters driving the dynamics of gynodioecy in natural population. Male fitness results confirm our previous studies: the D mitotype is associated with male sterility [41] while the K mitotype individuals are male-fertiles (restored hermaphrodites; [47]) but the penetrance of CMS does not appear to be total. Salient results are that temperature did not restore male sterility of D individuals or reverse male fertility of K mitotype. Moreover, temperature-dependent female advantage and costs, mediated by growth, were measured for D male-steriles (D > N normal hermaphrodites) and K restored hermaphrodites (K < N), respectively. Below we discuss in detail these results and their potential implication for dynamics of gynodioecy.

### A Temperature effect on male status (sterile / fertile)

As expected, incompatibility between nuclear and divergent mitochondrial genomes D and K induced a conflict over reproduction in the hermaphroditic snail *P. acuta*. Our observations confirm our previous studies on the male status of the three *P. acuta* mitotypes [41,47]. D individuals were almost all male-steriles (*i.e.,* they were unable to fertilize a virgin partner). The inability of D snails to fertilize a partner was associated with a smaller seminal vesicle containing few or no spermatozoa inside. David et al [41] associated these results with a reduction of male reproductive behaviour for D snails. However, as already noticed by David et al [41], the penetrance of CMS genes on male sterility is not total in D individuals. Here, a few of them (between 8% to 12%) managed to fertilize their partner. These individuals had small seminal vesicle size as did the other D individuals, but they sometimes produced more sperm, though always less than N or K individuals (see supplementary Fig. S6 and S7). This suggests that D does not fully wipe out sperm production, and that the remaining sperm sometimes passes the threshold necessary to fertilize some eggs of a virgin partner in the absence of competition. In gynodioecious plants, quantitative variation in male sterility phenotypes has also been observed [53–55], and often attributed to the presence of a polygenic or quantitative restoration system. In another context of reproductive manipulation, incomplete penetrance of parthenogenesis phenotype has been reported for the bacterial symbiont *Wolbachia* where infected females occasionally produced males [56]. However, in all these systems, it is difficult to tease apart cytoplasmic, nuclear and nongenetic variation on residual male fertility.

In contrast to D, almost all K snails were male-fertiles (*i.e.,* they were able to fertilize a virgin partner). Laugier et al [47] showed that introgression of the K mitochondrial lineage into a “naïve” nuclear genome induced a reduction of male fertility, which indicates that most K individuals, in their native genomic background, bear nuclear genes that restore male fertility. This restoration is very efficient as we found no aspect of male function (fertility, seminal vesicle and sperm) that differed significantly between K and N so far. In our study, although some K individuals did not fertilize their partner (between 4% and 8%), the proportion of failures was the same as in the N individuals (between 6% to 12%). Most of these N and K individuals that failed to inseminate their partner nevertheless had normal seminal vesicle size and sperm production, suggesting that they represent accidental failures rather than true long-term male sterility of the individual (see supplementary Fig. S6 and S7). A few exceptions of K individuals with some characteristics of a D-like phenotype (two K individuals with very low sperm counts at 18°C, 6 K individuals with relatively small seminal vesicles for their size; see Fig. 4 and Fig. S4) are compatible with incomplete fixation or penetrance of restorer genes in the K population, and overall their frequency was insufficient to draw inferences on the nature of this variation.

Contrary to observations in plant species where temperature changes (up or down) may restore fertility in male-steriles and reverse male fertility in restored hermaphrodites [25,26,28–33], the temperature did not interact with the male status of *P. acuta* mitotypes: D snails stayed male-sterile and K male-fertile at both 18 and 25°C. Two hypotheses may explain our results. First, the influence of environmental conditions on CMS and nuclear restorer genes may be dependent on genotypes (*i.e.,* genotype x environment interaction). For example, male sterility in barley or male fertility restoration in chives were influenced by temperature only in particular lineages [31–34]. This may suggest that CMS and restorer genes in the *P. acuta* populations studied here were not sensitive to temperature. Second, the temperatures tested, in the range of *P. acuta* tolerance, may not have been extreme enough to influence on male sterility/fertility at the life stage (*i.e*., adult) at which it was tested here.

### B Metabolism

Mito-nuclear incompatibility is known to interfere with mitochondrial function which can lead to a reduced fitness or fertility [57,58]. In plants, the mitochondrial genome divergence may be associated with an alteration of mitochondrial functioning (OXPHOS) and seems to cause male sterility in CMS individuals [10]. Surprisingly, despite the extreme genomic divergence of D and K mitotypes relative to N ones, we did not detect any difference in *in vivo* metabolic rate, between D, K and N mitotypes regardless the temperature. This can suggest that some mechanisms in D and K may compensate at the individual level for a dysfunction of the mitochondrial function (*e.g.* higher density of mitochondria in cells or particular OXPHOS functioning; [59]). In K restored hermaphrodites, the nuclear restorer genes could also influence the OXPHOS functioning [60]. Such mechanisms remain to be investigated in detail in gynodioecious *P. acuta*.

### C Temperature effect on fitness key traits involved in gynodioecy

Gynodioecy is a common reproductive system in plants which is defined by the coexistence of male-steriles (*i.e*., females) and hermaphrodites maintained through a female advantage in male-sterile individuals (in balance with a cost of bearing CMS genes) and costs (CMS and/or restoration) for restored hermaphrodites [19]. Taking these key parameters into consideration is essential and requires integrating how the environment modulates them to study the evolutionary dynamics of gynodioecy in natural populations.

#### C1 Temperature effect on FA: comparison between D male-steriles and N normal hermaphrodites

The FA in male-sterile individuals is often explained as an energy reallocation of male resources (saved from the stopped pollen production) toward growth and/or female function [6,61] or whole plant [18]. Eckhart [62] showed in *Phacelia linearis* that male steriles had a higher shoot biomass and seed production (at any shoot biomass) than hermaphrodites. Poot [61] found similar results in *Plantago lanceolata* where male-steriles had a higher dry mass and reproduction than hermaphrodites. Our results confirmed these theoretical predictions and empirical observations [16,18], demonstrating a female advantage in *P. acuta* gynodioecious species based on an energy reallocation from defective male function in D male-steriles toward body mass and a cascading effect on egg production. Indeed, D male-sterile snails globally had a higher growth rate relative to N normal hermaphrodites. This higher growth in D snails allowed them to increase their female fitness (i.e., higher egg production compared to N hermaphrodites) by approximately 20%. FA was influenced by the temperature as D individuals have a higher body mass than N, but this difference was only statistically significant at 18°C but not at 25°C. It is not surprising as growth is well known to be thermosensitive in *P. acuta* [50]. Contrary to body mass, shell length and the number of eggs were not influenced by temperature. In summary, the FA in D *P. acuta* male-steriles was driven by growth and this growth advantage impacted fecundity and was temperature-dependent. Our finding of a significant female advantage of the D mitotype seems to contradict the initial study of David et al [41], that failed to detect it. However, these authors worked at 25°C (where the FA is lower) and still found a (non-significant) 10% difference in egg production between D and N, with a very large variance and smaller sample size. The two studies are therefore qualitatively concordant, although the present one has more statistical power.

This is consistent with the maintenance of D male-steriles in gynodioecious populations and allows to predict that CMS could spread with an increase in frequency of D mitotype but depending on the thermal environment: it would be more likely to invade in cold conditions, as it can be less optimal for *P. acuta*. Correlations between environment and female (male-sterile) frequency are common in plants gynodioecious species, where more female (male-sterile) are found in harsh condition [40]. However, in contrast to our results, Bailey et al [36] did not find that temperature affect the magnitude of FA in *Lobelia siphilitica* although male-steriles individuals were more frequent in population where annual mean temperature are higher.

#### C2 Temperature effect on costs: K restored hermaphrodites vs N normal hermaphrodites

The model proposed by Dufaÿ et al [19] mentioned two potential costs to explain the maintenance of gynodioecious populations. Firstly, individuals who carry CMS sterilizing genes may suffer from a cost acting on the female function compared to non-CMS hermaphrodites (*i.e.,* normal hermaphrodites) named the cost of CMS. This CMS cost explains the maintenance of both non-CMS (normal) and restored hermaphrodites in gynodioecious populations. For example, in *Plantago maritima,* females that benefited from a higher number of seeds (FA) produced seeds with a lower germination rate (CMS cost) than hermaphrodites [22]. Secondly, individuals who have restorer genes also suffer from a cost (restoration cost) that may affect male and/or female fitness [6,13,19]. Some studies showed male fitness cost through low pollen viability in hermaphrodites carrying restorer genes (restored hermaphrodites) in *Lobelia siphilitica* [23] and in *Beta vulgaris* ssp *maritima* [54]. This cost of restoration may also affect female fitness as in *Plantago lanceolata* (reduced seed biomass; [63]) and *Phacelia dubia* (reduced seed viability; [64]), or both male and female fitness as observed in *Thymus vulgaris* [65]. However, both costs (of CMS and restoration) are intertwined which makes it challenging to differentiate them as they can act on the same fitness traits. In our study, K restored hermaphrodites and N normal hermaphrodites were very similar in terms of male fitness traits but K showed a slower growth rate than N. Moreover, this growth rate difference was more pronounced at 18°C than at 25°C (significant only at 18°C for body mass but at both temperature for shell length), suggesting a temperature-dependent cost for this mitotype. The difference in body mass translated into a slight disadvantage of K individuals in terms of female fertility at 18°C (−9%) but the latter was not significant.

Differences between N and K (and especially, the growth disadvantage in K) in our experiment must be interpreted in light of the high frequency of restorers in the common population they were extracted from, attested by the rarity of male-sterile phenotypes in the K category. Within the population, the two mitotypes coexist and exchange genes at each generation so we expect that both N and K mitotypes carry restorers in their nuclear genome, although the restorers are “silent” in the N context. This is attested by Laugier & Skarlou (in prep) who showed that offspring of male-sterile introgressed K individuals (in which the nuclear background carries no restorer) often become male-fertile again when their father comes from the laboratory N population from Lyon (that we used here as source of N snails). One condition for the maintenance of gynodioecy is that in a context of high restoration, the female fitness of CMS-bearing individuals (here, K) should be lower than that of non-CMS individuals (here, N), representing a cost of CMS in the restored context, otherwise nothing prevents the joint fixation of CMS and restoration in the population (reverting to 100% hermaphrodites [19]). The difference in growth observed here seems to fit with this condition. Laugier et al (submitted) recently observed than in captive evolving populations founded by mixing the N and K Lyon populations used in the present experiment, the frequency of K rapidly decreased over generations, indicative of a very large female fitness disadvantage of K (−18%). They also observed that the survival from egg to juvenile stage was slightly lower than that of N. Although neither the difference in egg survival (around −5%, Laugier et al. submitted) nor the difference in female fertility observed (around 9%, this study) seem sufficient by themselves to explain a female fitness difference of 18%, their combination does seem to be in the right range; and fitness differences may be enhanced by competition in the very dense (150 snails per aquarium) experimentally-evolving population of Laugier et al (submitted) compared to the isolated condition in which traits were measured. Strikingly, a common characteristic of growth differences (observed here) and egg-survival differences (observed previously) is to be enhanced at colder temperatures, suggesting again that the cost of CMS might be temperature-dependent.

## CONCLUSION

In the *P. acuta* gynodioecious system, we showed that D male steriles have a female advantage over N normal hermaphrodites and that, in a context of high restoration, K restored hermaphrodites have a cost associated to a lower growth than N. This pattern is consistent with the theoretical predictions for stable gynodioecious systems [19]. Female advantage and cost of CMS are key parameters involved in gynodioecy dynamics as they may change the frequencies of D male-steriles and K restored hermaphrodites among populations; in particular the frequency-dependence of female fitness, *i.e.* that female advantage of CMS predominates in a non-restored context, while female costs predominate in a restored context, is necessary to maintain gynodioecy. Our results lend support to this context-dependent expression of benefits and costs of CMS in the snail system. Moreover, although male sterility did not seem influenced by temperature, our study showed that environment may impact its benefits and costs in terms of growth and female fertility, which are higher at low temperature (18°C) than at 25°C (species optimum). Body mass seems to play a central role as our results show that female advantage and costs are mostly mass related. Thus, the dynamics of gynodioecy in *P. acuta* in nature may be affected by temperature. Other environmental factors like biotic interactions (*e.g*., predation) and intrinsic characteristics of populations (*e.g.,* population size, inbreeding, connectivity; [20]) could also influence relative fitness and ultimately gynodioecy dynamics. Consequently, inferring evolutionary dynamics of gynodioecious populations from laboratory measurements of fitness traits may be very challenging, and should be completed by direct tracking of temporal dynamics of CMS and restorer genes in wild populations.

## Acknowledgements

We are grateful to Loïc Trebignaud who helped in snail dissections.

## Author contributions

P.D., S.P. and E.L. conceived the project. S.B., S.P. and E.L. performed the experiment. L.T. performed the *in vivo* metabolism measures. S.B. analysed the data and wrote the first version of the manuscript. All author revised the manuscript.

