## Supplementary material for "Temperature-dependant benefits and costs of cytoplasmic male sterility in snail *Physa acuta*"

| Population | Mitotypes frequencies | GPS coordinates |
|---|---|---|
| Autoroute | 14%D, 86%N (2015, 2016,2017) | N45°48'05.3", E 4°55'26.1" |
| Erevan | 68%N, 32%K (2019) | N 45°74'42.1" E 4°81'58.4" |
| Irigny | 85%N, 3%D, 12%K (2019) | N 45°68'20.1" E4°83'38.7" |

Table S1. Mitotype frequencies and GPS coordinates of source populations of the three mitotypes

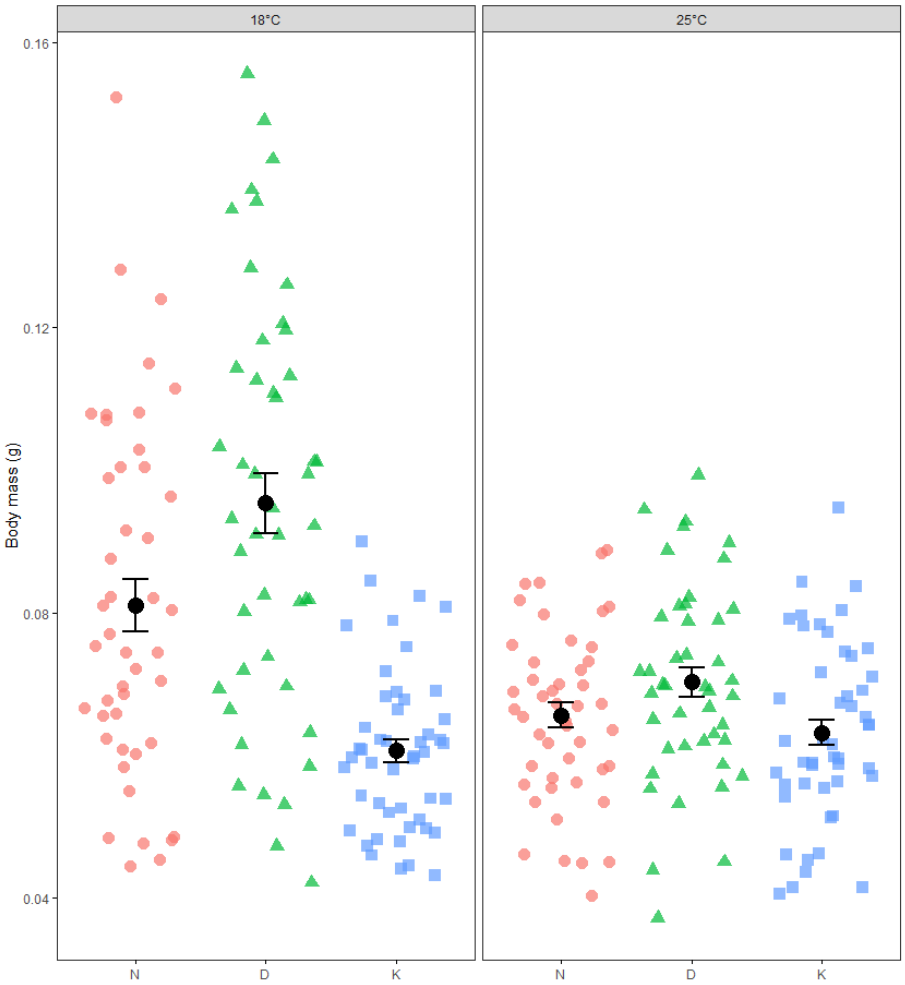


Figure S2. Body mass for normal *P. acuta* hermaphrodites N (red circle), male-sterile D (green triangle) and restored hermaphrodites K (blue square) at 18 and 25°C. Body mass was measured at 45 and 22 days for 18°C and 25°C treatments, respectively. Dots correspond to

observed values. At 18°C, the mean body mass was 0.0810 ± 0.0037 g for N, 0.0953 ± 0.0043 g for D and 0.0606 ± 0.0016 g for K individuals. At 25°C, the mean BM was 0.0656 ± 0.0017 g for N, 0.0703 ± 0.0021 g for D and 0.0632 ± 0.0018 g for K individuals.

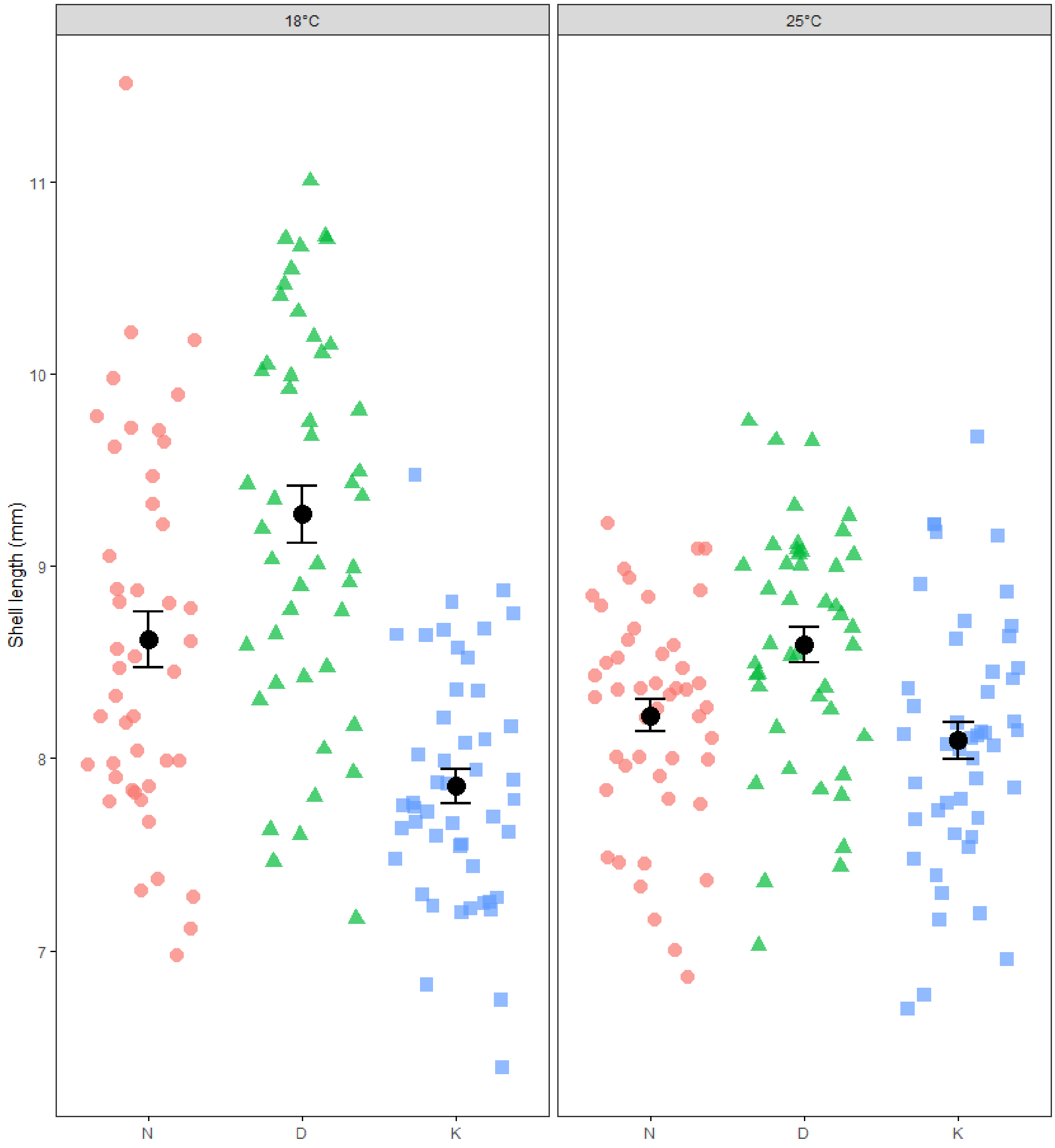


Figure S3. Shell length for normal P. acuta hermaphrodites N (red - circle), male-sterile D (green - triangle) and restored hermaphrodites K (blue square) at 18 and 25°C. Length was measured at 45 and 22 days for 18°C and 25°C treatments, respectively. Dots correspond to observed values. At 18°C, the mean shell length was 8.62 ± 0.98 mm for N, 9.27 ± 1.02 mm for D and 7.85 ± 0.62 mm for K individuals. At 25°C, the mean shell length was 8.22 ± 0.57 mm for N, 8.59 ± 0.63 mm for D and 8.09 ± 0.66 mm for K individuals.

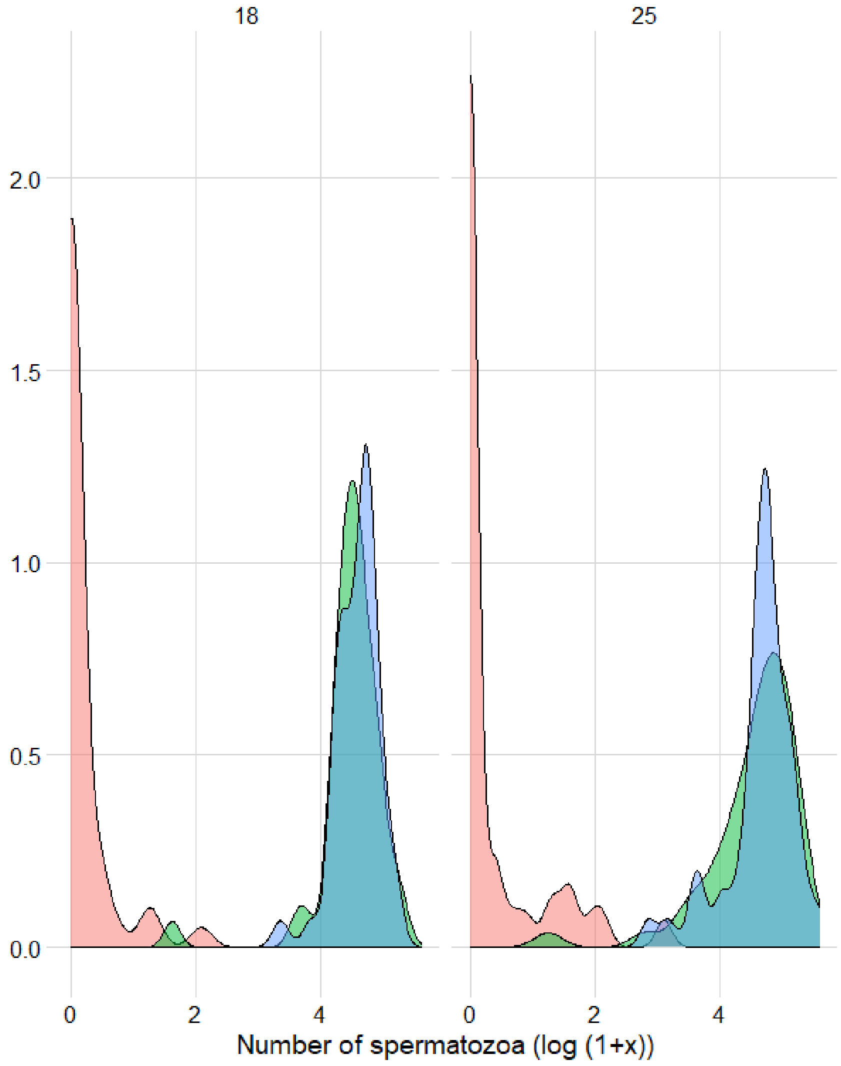


Figure S4. Distributions of sperm counts (means between two individual counts) in adults of normal hermaphrodites N (red), male-sterile D (green) and restored hermaphrodites K (blue) at 18 and 25°C.

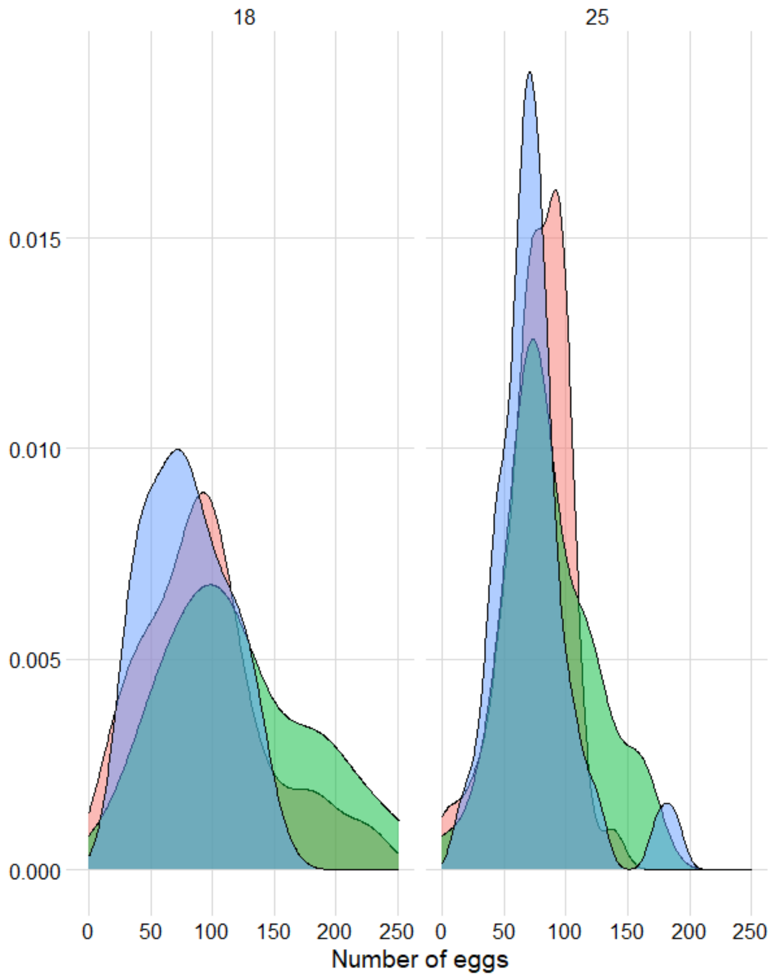


Figure S5. Distributions of egg number (female fitness) produced by focal individuals of normal hermaphrodites N (red), male-sterile D (green) and restored hermaphrodites K (blue) at 18°C and 25°C.

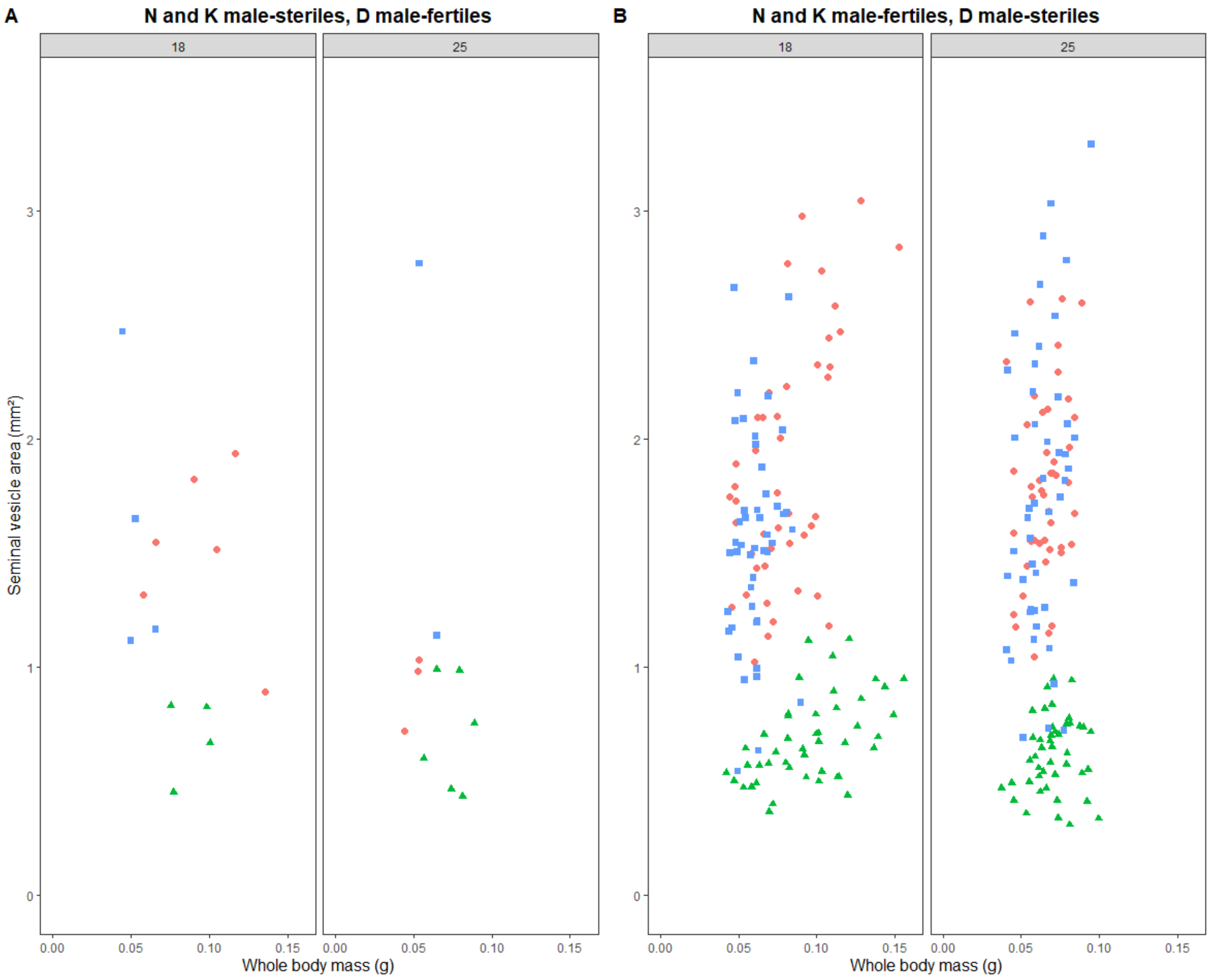


Figure S6. Seminal vesicle area (male fitness) according to body mass for normal *P. acuta* hermaphrodites N (red - circle), male-sterile D (green - triangle) and restored hermaphrodites K (blue - square) at 18°C and 25°C. **A –** N and K male-steriles and D male-fertiles. **B –** N and K male-fertiles and D male-steriles.

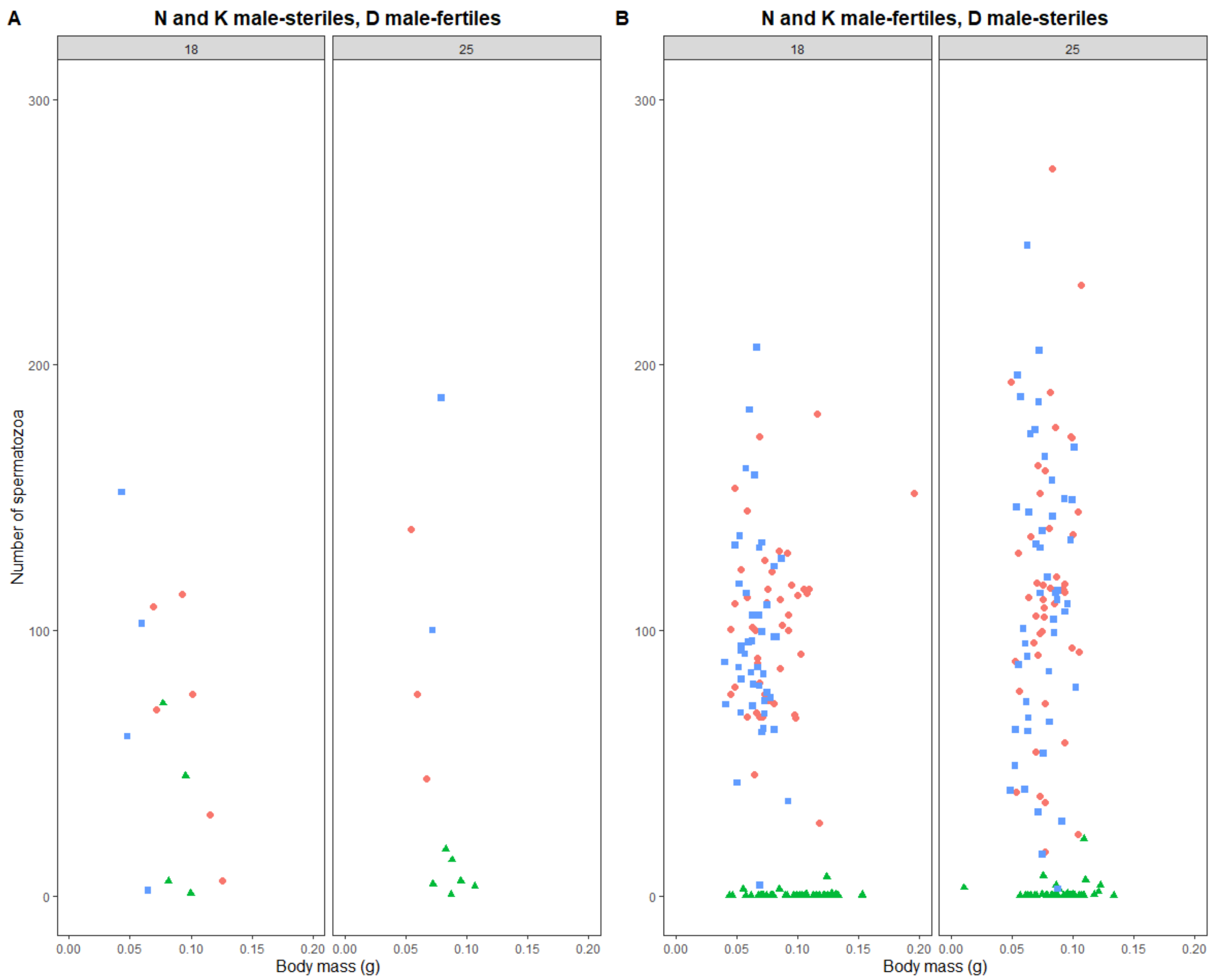


<u>Figure S7.</u> Number of spermatozoa (means between two individual counts for a sake of clarity) according to body mass for normal *P. acuta* hermaphrodites N (red - round), male-sterile D (green - triangle) and restored hermaphrodites K (blue - square) at 18°C and 25°C. **A –** N and K male-steriles, D male-fertiles. **B –** N and K male-fertiles, D male-steriles.
